# Mining for ions: diagnostic feature detection in MS/MS spectra of post-translationally modified peptides

**DOI:** 10.1101/2022.09.12.507594

**Authors:** Daniel J. Geiszler, Daniel A. Polasky, Fengchao Yu, Alexey I. Nesvizhskii

## Abstract

Post-translational modifications (PTMs) are an area of great interest in proteomics, with a surge in methods to detect them in recent years. However, PTMs can introduce complexity into proteomics searches by fragmenting in unexpected ways. Detecting post-translational modifications in mass spectrometry-based proteomics traditionally relies on identifying ions shifted by the masses of the modifications. This presents challenges for many PTMs. Labile PTMs lose part of their modification mass during fragmentation, rendering shifted fragment ions unidentifiable, and isobaric PTMs are indistinguishable by mass, requiring other diagnostic ions for disambiguation. Furthermore, even modifications that have undergone extensive characterization often produce different fragmentation patterns across instruments and conditions. To address these deficiencies and facilitate the next generation of PTM identification, we have developed a method to automatically find diagnostic spectral features for any PTM, allowing subsequent searches to take advantage of additional metrics and increase PTM identification and localization rates. The method has been incorporated into the open-search annotation tool PTM-Shepherd and the FragPipe computational platform.

## Introduction

Post-translational modifications (PTMs) have long been of interest to proteomics researchers because of their central role in regulating cellular functions. Processes to maximize their recovery run the gamut of proteomics techniques, from sample preparation^1^ to instrumental acquisition^2^ and computational analysis^3-5^. At the computational level, proteomics search engines have grown tremendously in their capacity to identify PTMs. For PTMs with complex fragmentation patterns like glycosylation that exhibit multiple modes of fragmentation, entire search engines specific to the modification class have been developed^4,6,7^. Despite this work, many modifications continue to suffer from low recall in standard high-throughput workflows due to their behavior during tandem mass spectrometry (MS) analysis, producing unexpected or difficult fragmentation patterns that frustrate search engines^8^. Even small changes to workflows—such as the addition of isobaric labels—can alter fragmentation patterns and reduce or preclude identification of even the best-studied PTMs^9^. Recent work with synthetic peptides carrying less well-studied PTMs demonstrated that many diagnostic ions and neutral losses have yet to be identified^10^.

With the proliferation of synthetic PTMs^11^—particularly ones that alter fragmentation patterns^9^— and new instrumental methods^2,12^, keeping search engines up to date with knowledge of how an analyte will fragment in a particular setting is a herculean task. To overcome this, computational tools are being developed to identify modification fragmentation patterns without prior knowledge. The first such tools only identified diagnostic ions and were limited in their applications^13^, but newer approaches have incorporated additional features. Synthetic peptides bearing modifications are generally seen as a gold standard to study PTM fragmentation patterns and methods have been developed to extract them from spectra^14^, but this approach adds additional benchwork to proteomics experiments. Furthermore, optimal search parameters are fragmentation-dependent and can change based on experimental settings, which requires reprocessing mass spectrometry data and reanalyzing fragmentation patterns for multiple experiment types. Zolg et al. (2018) developed a method to do this in a high-throughput manner, but it requires paired modified-unmodified peptides and cannot be easily reimplemented by other research groups. Their approach to identify neutral losses also requires both the intact and fragmented peak to be present in the spectrum at consistent distances, precluding finding complete losses and many charged losses. Other approaches to score PSMs from modified peptides are trained for specific PTMs^15^ or perform model refinement that focuses on distances between experimental peaks, discarding information about matched ions from the peptide backbone that would dramatically reduce the required training dataset size^16^. Chemoproteomics has a particular stake in this effort due to the diversity of probes employed^17-19^. However, existing tools for chemoproteomics require isotopic labeling signatures to be present at the MS1 and MS2 levels^20^. This limits their applications to chemical probes that are labeled non-isobarically, thus they cannot be used for some PTM probes^21^, biological PTMs, or the development of isobaric mass tags^22^. In prior work, we have found that understanding PTM fragmentation patterns allowed us to maximize modified peptide recovery and localization. Thus, when studying a cysteine chemoproteomic probe, we developed a method to extract its diagnostic spectral features to improve coverage of the ligandable cysteineome^19^. Our approach did not require synthesizing standards or isotopically labeled peptides and facilitated the discovery of partial modification losses and diagnostic ions, ultimately leading to the identification of three diagnostic ions and two partial PTM fragmentation events that escaped manual inspection.

Here, we present an improved, fully automated, and empirically tuned implementation of our diagnostic feature extraction algorithm to study and score the fragmentation patterns of modifications. Our approach detects three separate types of diagnostic features—diagnostic ions, peptide remainder masses, and fragment remainder masses—and can be used in any experimental setting, including for the simultaneous characterization of multiple modifications and when only a handful of PSMs are present for a modification. We demonstrate the robustness of our technique by applying it at both massive and small scale, and across synthetic and biological PTMs. Finally, we perform a meta-analysis of diagnostic features and discuss how these can be used to further PTM discovery in diverse settings. Our method has been implemented within PTM-Shepherd^23^ and is freely available as part of the FragPipe suite of tools (https://fragpipe.nesvilab.org/).

## Results

### Algorithm overview

The PTM-Shepherd diagnostic feature mining module aims to perform high throughput identification of spectral features that can be used to identify post-translational modifications (PTMs), facilitating the validation or discovery of PTM-specific signals. Probable modifications from an experiment are identified by passing the results of open or mass offset search to PTM-Shepherd. For each MS1 mass shift, PTM-Shepherd identifies enriched diagnostic features across three categories: diagnostic ions; mass shifts from the unmodified, intact peptide ions (peptide remainder masses); and mass shifts from unmodified fragment ions (fragment remainder masses). This module operates in three steps: calculating all possible spectral features for every peptide-spectrum match (PSM) with a particular mass shift, identifying the most abundant spectral features for every identified mass shift within each category, then finally performing statistical tests and filtering to see whether those features can be used to infer the presence of the modification via comparison to unmodified peptides. This module uses as input decharged and deisotoped MGF spectra produced by MSFragger^24^, so the maximum charge state for all ions in MS/MS spectra is assumed to be one. Spectral ions are normalized to the base peak and only the top 150 peaks are considered (by default).

We illustrate our technique using a cysteine-specific chemical probe^19^ that we previously analyzed with an early version of the algorithm, identifying all three ion types (**Fig 1a**). The first step in our strategy is to calculate all possible diagnostic spectral features for each PSM within a mass shift identified by PTM-Shepherd. Any ions from experimental spectra that do not belong to the peptide are considered potential diagnostic features for the mass shift. To identify recurring features for the mass shift, calculated features for every spectrum from the mass shift are sent to a common histogram. Peaks are identified from here and shuttled to downstream analysis. For diagnostic ions, the unannotated ions from the experimental spectrum are sent to their histogram as they are (**Fig 1a**, green). Peptide remainder masses are calculated by computing mass differences between the theoretical, unshifted peptide ion (purple) and all ions in the spectrum (blue). Fragment remainder masses are calculated by iteratively computing mass differences between every theoretical ion from the peptide backbone (purple) and all ions in the spectrum.

**Figure 1:**
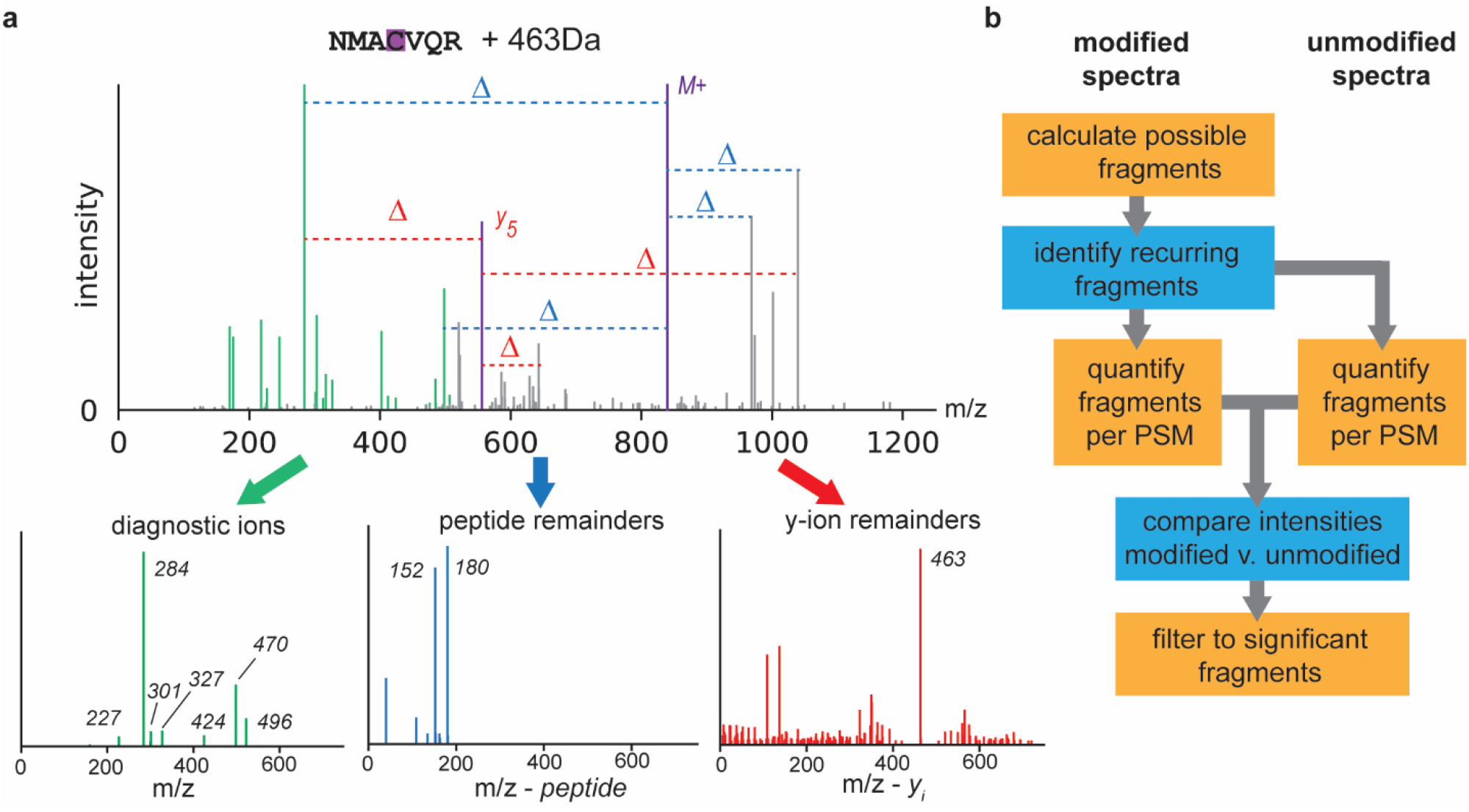
Implementation of the diagnostic feature mining algorithm. (a) Calculation of all possible diagnostic ions (green), with peptide remainders (blue) and fragment remainders (red) calculated with respect to theoretical unmodified ions (purple) for a synthetic Cys probe of mass 463 Da. Calculated features across spectra are subsequently pooled to find recurring features. (b) Workflow for diagnostic feature selection.

Finding recurring ions does not mean that they are useful for identifying a mass shift. Our ion set contains features that might be abundant across the entire dataset, so it is necessary to remove baseline noise. We do this by comparing the recovered features from all spectra bearing the mass shift to those of unmodified peptides in bulk as a proxy for dataset background (**Fig 1b**). For every feature detected in the prior step, it is quantified across modified and unmodified PSMs, with missing ions or offsets encoded as zeroes. The result is two lists of intensities, from which we can perform statistical tests. Encoding missing ions as zeroes is necessary for this step, but it can also produce a range of non-normal distributions, calling for the non-parametric Mann-Whitney-U test. Features that are significantly different between the modified and unmodified lists are then filtered for sensitivity criteria (minimum prevalence in the modified bin) and mean intensity fold-change between the two bins. Fragment remainder ions undergo an additional layer of filtering for ion formation propensity, where they are required to represent a minimum percentage of the number of ions in their series. True fragment remainder ions can also create “echoes” of their masses that are combinations of the original mass and adjacent amino acids, multiple of which can pass filtering for a mass shift. We correct these by checking for enrichment of adjacent amino acids from the residues the remainder mass is derived from and adjusting the mass accordingly. Because the adjacent residues are pseudo-random in most cases, we also reasoned that any fragment remainder mass less intense than the first corrected mass is likely to be noise. These are also filtered from the result. Additional details about this process can be found in the *Methods* section.

Re-analyzing the cysteine probe data used for illustration above, we identified all eight diagnostic features—five diagnostic ions (**Fig 1a**, green), two peptide remainder masses (**Fig 1a**, blue), and one fragment remainder masses (**Fig 1a**, red)—that were annotated in the prior study and are high confidence identifications. Furthermore, we also identified two additional diagnostic ions, suggesting improved sensitivity for the empirically tuned and automated algorithm (the full attribute list can be found at **Supplementary Table 1**).

**Table 1:** Most prevalent mass shifts and their characteristics from an open search of a pRBS-ID experiment. Two unannotated mass shifts were determined to be the most likely candidates for RNA moieties. This was confirmed by their similar effects on retention time.

| peak apex | PSMs | % also in unmodified | mass annotation | ret. time shift | localized PSMs | n-term. localization rate | AA1 | AA1 enrichment score | AA1 PSM count |
| --- | --- | --- | --- | --- | --- | --- | --- | --- | --- |
| 0.0002 | 7751 | 100 |  | -0.6 | 291 |  |  |  |  |
| 226.0594 | 4241 | 34.81 | 226.0594 mass-shift | -449.3 | 1343 |  | R | 2.0 | 160 |
| 1.0032 | 2378 | 88.62 | +1 isotopic peak | 25.6 | 658 |  | F | 1.6 | 65 |
| 57.0216 | 945 | 80 | Iodoacetamide derivative | -63.8 | 806 | 38.94 | H | 11.9 | 184 |
| 94.0168 | 830 | 50.81 | 94.0168 mass-shift | -477.6 | 799 | 7.47 | H | 9.2 | 140 |
| -48.0032 | 568 | 35.35 | Homoserine lactone | -1238.7 | 559 | 26.23 | M | 53.0 | 373 |

### New protocol characterization

RNA-crosslinking studies also feature labile modifications. Repeating sugar molecules can fragment in myriad ways, frustrating attempts to localize or even identify RNA moieties. Bae et al. recently developed pRBS-ID, an RNA crosslinking workflow utilizing photoactivatable nucleotides and chemical RNA cleavage to overcome these challenges^25^. Alongside the development of their bench technique, they needed to develop a bespoke computational workflow to identify RNA fragment remainder masses and identify and quantify their host peptides. We believed that this process could be recapitulated by PTM-Shepherd without the need for time-intensive custom workflows, and as such we struck a course to replicate their results for the commonly used 4-thiouridine (4SU) nucleotide analog^26^.

First, we performed an open search using the default diagnostic ion mining setting available in FragPipe. As expected in any open search, PTM-Shepherd identified many mass shifts for biological and chemical PTMs, but two unannotated mass shifts of 226 Da and 94 Da at high amounts likely corresponding to the modification of interest (**Table 1**, full mass shift profile can be found at **Supplementary Data 1a**). These mass shifts matched those identified by Bae et al. Notably, the fragment remainder masses PTM-Shepherd identified for both mass shifts were nearly identical, indicating with a high degree of likelihood that they had the same source. In this case, fragment remainder masses of 94 Da were identified from both mass shifts’ *b*- and *y*-ion series, and an additional fragment remainder mass of 77 Da (the prior remainder with a loss of ammonia) was identified from both mass shifts’ *b*-ions (**Supplementary Data 1b**). Like the loss of ammonia described from glycopeptide’s *Y*-ion series, this mass shift appeared to be diagnostic for RNA-crosslinked peptides (226 mass shift AUC = 0.57, 94 mass shift AUC = 0.58, **Supplementary Note 1**).

After a more targeted search using these fragment remainder masses, we also wondered whether any additional diagnostic features might appear for the RNA-crosslinked peptides and performed a second pass at diagnostic feature mining (**Supplementary Table 2**). For peptide remainder masses, we recovered the two masses described above from both *b-* and *y-*ions (**Fig 2a,b**) as well as an *a-*ion associated mass shift at 66 Da (94 Da minus 28 Da) from *b*-ions. Diagnostic ions can be of particular interest for future analyses, such as in ion-triggered instrument routines, even if they are left unused at the present. We found two easily explicable diagnostic ions for the intact nucleoside (**Fig 2b**): an ion at 133 *m/z* corresponding to a dissociated ribose, the other half of the 94 Da fragment remainder mass, and an associated neutral loss of water. Accordingly, these ions were not diagnostic for the MS1 mass shift corresponding the nucleoside without the ribose (**Fig 2d**), as with the ribose already dissociated there is nothing left to form the diagnostic ion.

**Table 2:**
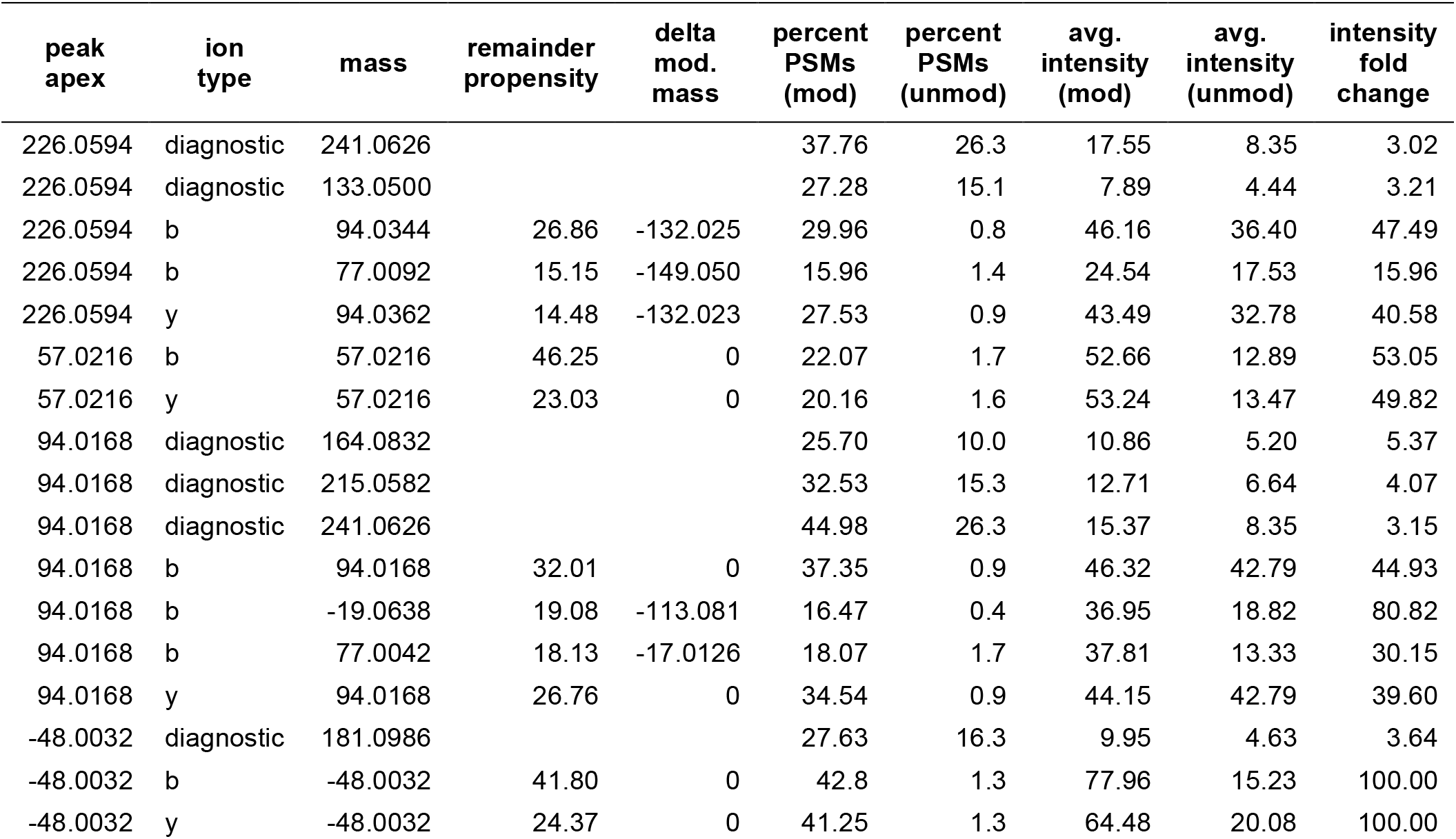
Diagnostic features for the most abundant mass shifts detected in an open search of a pRBS-ID experiment. Remainder propensity scores are present only for *b-* and *y-* remainder masses. Features corresponding to the set of modified or unmodified PSMs used in comparisons are labeled as (mod) and (unmod), respectively. The difference between the MS1 mass shift and the observed fragment remainder masses (i.e., the lost mass) is enumerated in the “delta mod mass” column.

**Figure 2:**
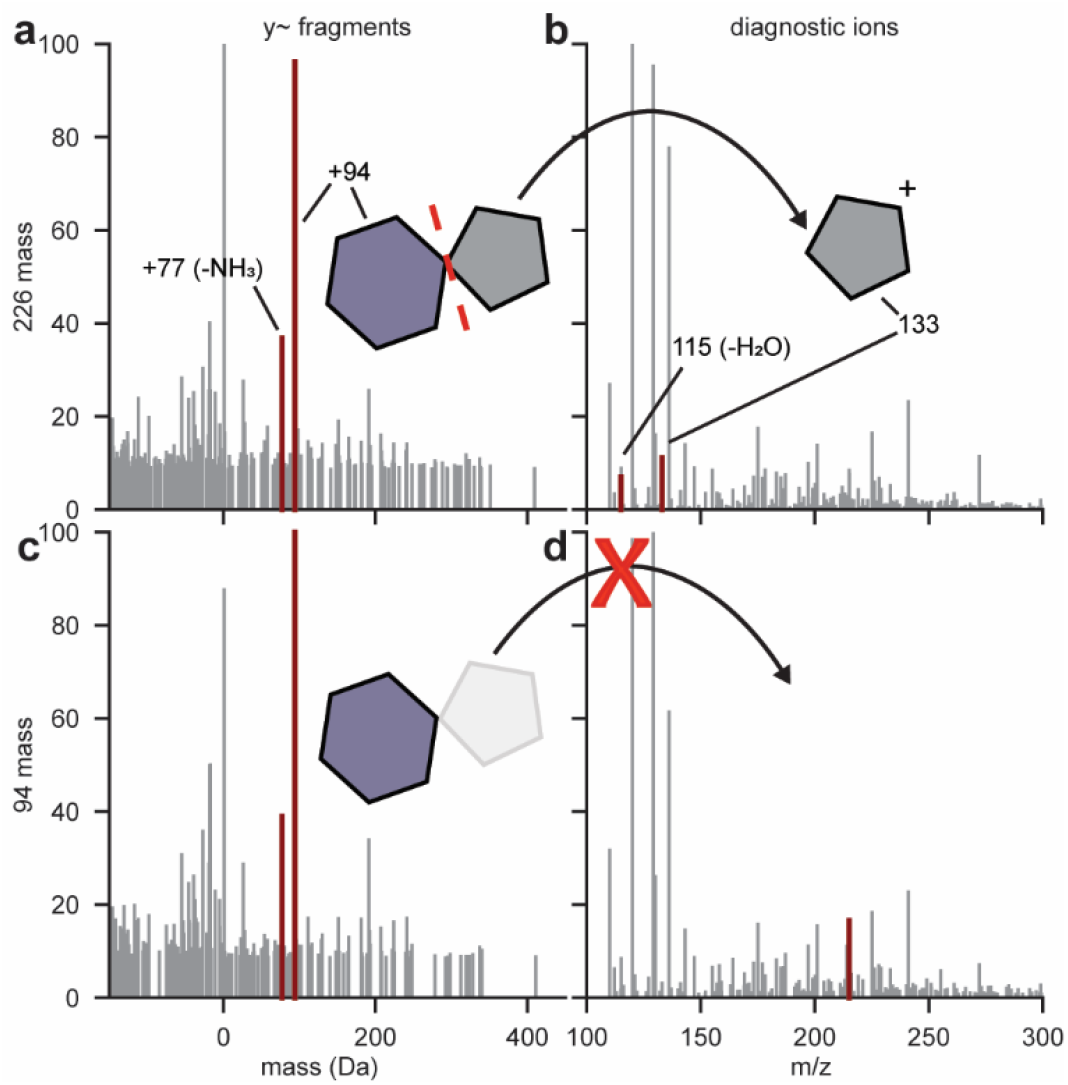
Characterization of 4SU fragmentation from a pRBS-ID experiment. (a) All possible fragment remainder masses for *y-*ions from the 226 Da mass shift. Remainder masses that passed PTM-Shepherd’s filtering are highlighted in dark red, corresponding to the retention of the 94 Da fragment on the peptide. Figure 3b-d use the same color scheme as Fig 3a. (b) Diagnostic ions derived from the fragmentation of the nucleoside analog from the 226 Da mass shift. (c) All possible remainder masses for *y-*ions from the 94 Da mass shift. (c) All possible diagnostic ions from the 94 Da mass shift.

### Data-driven discovery of diagnostic features

Glycopeptides contain labile modifications that produce rich sequences of diagnostic ions and peptide and fragment remainder masses^27^. We reasoned that detecting known glycopeptide fragmentation patterns would be a good way to validate our algorithm’s performance given the extensive literature characterizing glycopeptide fragmentation. To this end, we searched for glycopeptides in a large IMAC-enriched, TMT-labeled clear cell Renal Cell Carcinoma (CCRCC) dataset^28^. Phosphorylation enrichment by IMAC, the method employed in this publication, has been shown to simultaneously enrich glycopeptides, particularly those bearing sialic acids^29,30^, so the data should be rich in glycan signals. This dataset also presents two challenges: TMT-labeling is known to affect PTM fragmentation patterns due to reduced proton mobility^9^ and the relatively high collision energies used in this experiment cause extensive fragmentation of glycans, reducing the signal strength of typical glycan fragment ions.

We first wanted to verify that we could detect the commonly used glycopeptide-associated diagnostic ions from the MSFragger-Glyco^7^ search and annotation that are most likely to be present^31^. After discarding any mass shifts less than 50 Da we were left with 493 likely glycan mass shifts from 967,264 glyco PSMs of 9623 unique glyco peptides, each of which should be enriched for diagnostic ions associated with the N-glycan core structure^12^ and other monosaccharide(s) present, including sialic acid. Indeed, PTM-Shepherd successfully identifies many of the expected diagnostic ions used in glycopeptide searches and glycan identification ^7,31^, including three known sialic-acid related oxonium ions at 274, 292, and 657 m/z (**Fig 3a, Supplementary Data 2**). In addition to these, we found 12 additional ions that were diagnostic for more than 50% of glycan mass shifts. We hypothesized that these might be diagnostic ions specific to a high-collision energy environment and attempted to identify them in a data-driven manner. We used PTM-Shepherd’s diagnostic feature extraction module, which extracts intensities for user-specified ions of interest, to quantify these alongside the set of common diagnostic ions used in the MSFragger-Glyco, identifying clusters of highly correlated ions (**Fig 3b**, see *Methods*). Known ions clustered together meaningfully, with annotated GalNac, Hex, HexNac, and PhosphoHex ions being highly correlated with others from the same residue, lending credence to this approach’s validity. Perhaps unsurprisingly given the nature of the enrichment method, most unannotated diagnostic ions formed a large cluster with the two monomeric sialic acid oxonium ions found at 274 and 292 *m/z*. We selected the diagnostic ions from a subcluster (**Fig 3b**, cluster 5) that was highly correlated with both oxonium ions (**Supplementary Data 3**) to validate individually. These ions formed a potential neutral loss series from the annotated 292 and 274 m/z oxonium ions, with successive losses of 42, 17, 18, and 30 Da. To our knowledge, manuscripts covering sialic acid fragmentation make no mention of these as diagnostic ions^12,32,33^, so their presence in spectra acquired at high collision energies may be of interest to other researchers when assigning sialic acids to glycan composition.

**Figure 3:**
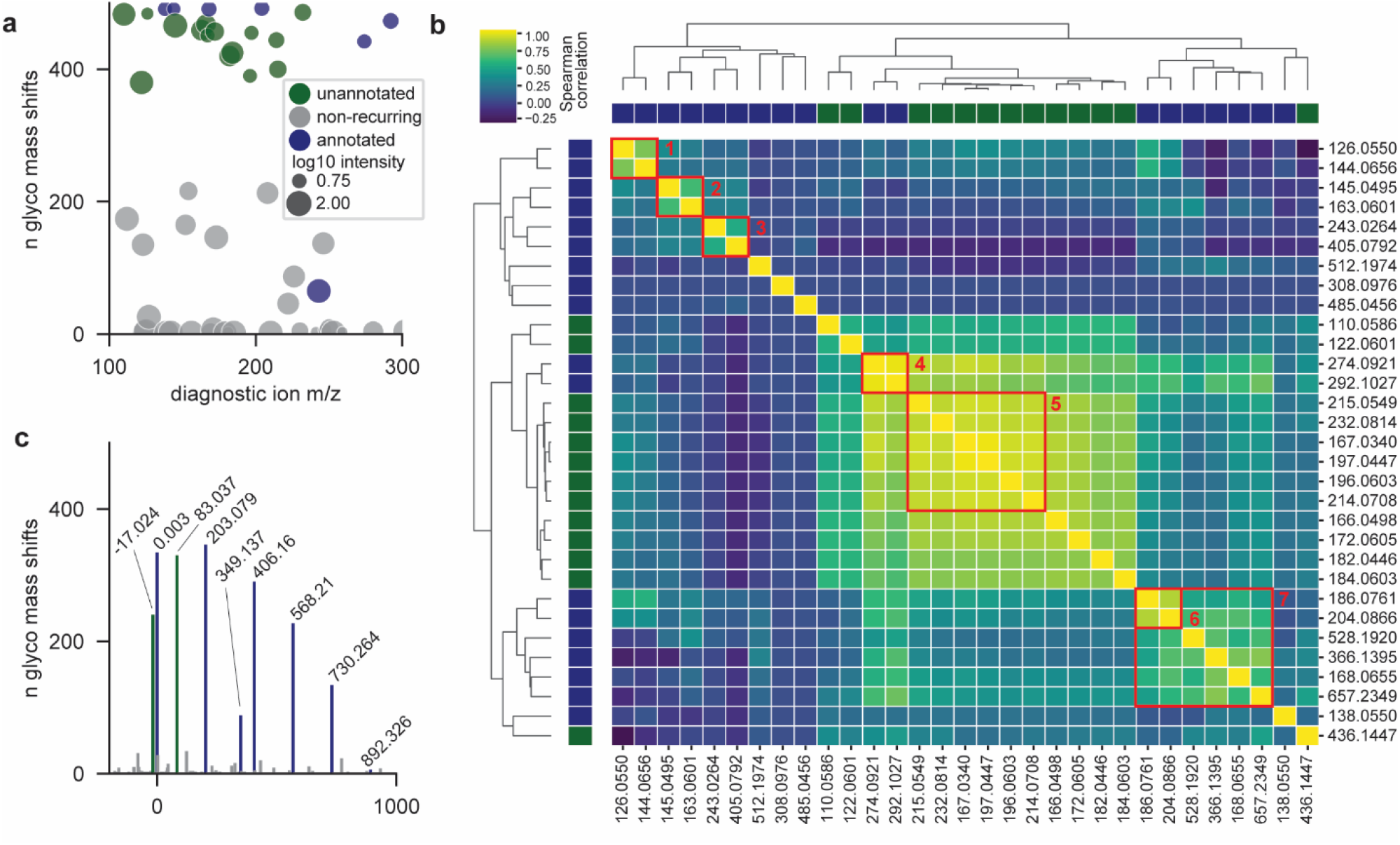
Diagnostic features of IMAC-enriched glycopeptides under high energy conditions. (a) Scatterplot of recovered diagnostic ions across glyco mass shifts. Ions occurring in >50% of mass shifts are considered recurring and are included in Fig 2b. Color schemes for Fig 2b-d are consistent with Fig 1a. (b) Spearman correlation clustering between diagnostic ions across all glyco spectra. Identifiable clusters are as follows: 1: GalNac, 2: Hex, 3: PhosphoHex, 4: NeuAc, 5: sialic acid, 6: HexNac monomers; 7: HexNac including non-monomers. (c) Histogram of peptide remainder ions across glyco mass shifts. (d) Histogram of fragment remainder *b-*ions across mass shifts.

Aside from diagnostic ions, glycopeptides also produce an intense series of peptide remainder ions, called *Y*-ions in glycopeptide fragmentation nomenclature, where the peptide is intact while the modification has fragmented^12^. Mammalian N-glycans have a common core structure. When the core structure fragments, it produces a pattern of *Y*-ions with peptide remainder masses that are identical irrespective of the peptide’s or glycan’s mass and can even be used to diagnose the presence of glycopeptides^6^. Like the diagnostic ions discussed above, we find an expected pattern of peptide reminder masses corresponding to the N-glycan’s core’s *Y*-ion series (**Fig 3c**). Aside from these, two peptide remainder masses that are not considered in the MSFragger-Glyco search recurred across mass shifts: +83 Da and -17 Da. The smallest glycan mass from the N-glycan core, corresponding to a single GlcNAc retained on the peptide, is 203 Da, so seeing masses smaller than that being as diagnostic for glycopeptides as the complete loss of glycan (+0 Da) or a single GlcNAc (+203 Da) was surprising. This pattern—consisting of a cross-ring fragmentation event at the core GlcNAc and a loss of an ammonium, respectively— has previously been identified as a conserved fragmentation pattern for glycopeptides^34^, but appears not to be used in current state-of-the-art tools^6,7,35^. This indicates that even for very well characterized modifications, gaps can exist between knowledge of fragmentation patterns and their use in computational tools, a disconnect that PTM-Shepherd’s automated fragmentation analysis can correct.

The final diagnostic feature we assessed for this glycan dataset is shifted fragment ion series. When the peptide and glycan have both fragmented, the glycan can leave a signature +203 fragment remainder mass on the peptide ion series^12^. PTM-Shepherd recovered this fragment remainder mass exactly (**Supplementary Figure 1**) and with little interference from artefactual mass shifts despite the noisy nature of pairwise ion differences.

Some of the identified ions, particularly the *Y*-ion series of peptide remainder masses, appeared to taper off very quickly at larger masses, which is a known issue when identifying labile modifications at relatively high collision energies. We reasoned that using these extra ions in our search when they can be low-abundance or absent injects additional noise into the search results and suppresses real glycopeptide identifications. To test this, we used the fragmentation information provided by PTM-Shepherd and reduced our fragment and peptide remainder masses to only the four *Y-*ions appearing in >50% of glycan mass shifts. Though more careful analysis would surely yield better results, even the incorporation of a crude cutoff from a subset of the data resulted in a 4.5% increase in glyco-PSMs, proving that the fragmentation information provided by PTM-Shepherd enables researchers to tune search parameters to best suit their individual experiments.

We showed that PTM-Shepherd was sensitive to known diagnostic features for glycopeptides. New features detected by PTM-Shepherd also had chemical meaning relevant to the experimental setting, and PTM-Shepherd was able to identify unannotated sialic acid diagnostic ions for high-energy TMT experiments in a data-driven manner. Additionally, we proved that the information provided by PTM-Shepherd can be incorporated into subsequent searches to increase to fine-tune parameters for different experimental settings.

### Automated fragmentation analysis

ADP-ribosylation (ADPR) has seen a surge of interest in recent years, with many enrichment methods^36,37^, and instrumental techniques^38^ developed over the last decade to aid in its study. Despite this, specialized computation techniques have lagged behind. Fragmentation studies— necessary to design tools or workflows for the analysis of PTMs—require painstaking analysis and examination of individual spectra^39^. We believed that PTM-Shepherd’s diagnostic feature mining module could expedite fragmentation studies and reveal new, useful insights to their behavior. To demonstrate this, we reanalyzed ADPR-enriched data from Martello et al.^38^ from peroxide-treated HeLa cells, rich in Ser-directed ADPR, and mouse liver, rich in Arg-directed ADPR.

To validate the fragmentation patterns we detected, we first cross-checked them against published ones^39^. As expected, we found previously annotated diagnostic ions (**Fig 4a, Supplementary Data 5a**,**b**) corresponding to almost every expected breakpoint on the ADPR side chain (**Fig 4a**). These were all found at relatively high levels among ADPRylated spectra (78-100%). Interestingly, the most intense of these ions—e.g., the adenine-derived ion at 136— was also found at high levels in unmodified spectra (73%), meaning its presence was not specific to PSMs with ADPR (**Supplementary Note 2**). This speaks to the robustness of PTM-Shepherd’s algorithm; even features whose presence alone is not specific to a particular mass shift can be recovered because our scoring and filtering utilizes intensity information. We also recovered additional ions that correspond to derivatives of annotated ions: an oxidized 428 *m/z* ion (+16 Da), a 348 *m/z* ion that has undergone a loss of water (−18 Da), and a 250 *m/z* ion that has undergone a loss of water (−18 Da). These ions were all far more specific to the ADPR PTM than their annotated counterparts and thus may be of interest to others studying ADPR. A final diagnostic ion of interest did not correspond to a common mass offset from an annotated ion. At 137.0458 *m/z*, we could not identify this ion as being a secondary product of any annotated ions. Its exact mass is strongly suggestive of a deamidation event occurring on the adenine ion at 136.0618 *m/z* (+0.9840 Da).

**Figure 4:**
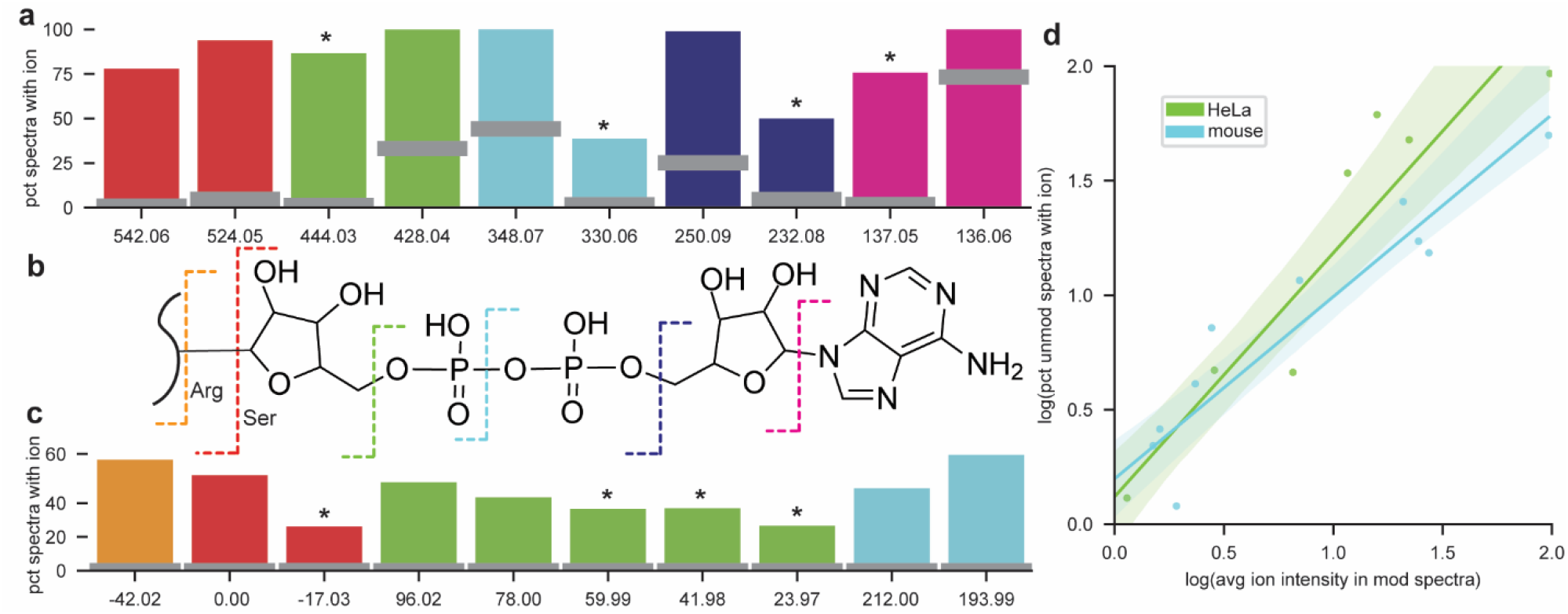
Analysis of ADPR fragmentation patterns. (a) ADPR diagnostic ions. Colored bars show the percentage of ADPR PSMs containing the diagnostic ion, while gray bars show the percentage of unmodified PSMs containing the diagnostic ion. Ions detected in both datasets were averaged across all values. A “*” denotes novel features discovered by PTM-Shepherd. (b) Structure of ADPR. Dashed lines correspond to breakpoints in the molecule, and color corresponds to the diagnostic features produced during fragmentation. (c) ADPR peptide remainder masses. (d) Correlation between average intensity of diagnostic ions and their presence in unmodified spectra across both datasets. Confidence intervals for Pearson’s correlation are highlighted.

We also observed a strikingly strong relationship between an ion’s average intensity and its presence in unmodified spectra across both ADPR datasets analyzed (**Fig 4c**, Spearman’s *R*^*2*^: mouse = 0.857; HeLa = 0.884). We have previously commented on this phenomenon when looking at biotin-derived Cysteine probes^19^. In that case, reducing the isolation window and employing ion mobility gave a modest boost to diagnostic ion specificity, an effect that was presumed to be caused by reduced co-fragmentation of peptides. It is worth noting that the issue of co-fragmented ions has been well-studied in the context of isobaric tandem mass tags^40^. But, to our knowledge, there has been little discussion of parallel issues when using diagnostic ions for biological PTMs.

PTM-Shepherd also identified both types of remainder ions in this dataset, peptide (**Fig 4d**) and fragment. Of note was PTM-Shepherd’s recovery of a -42 Da peptide remainder mass from the Arg-directed ADPR dataset (**Supplementary Data 5b**). When Arg-linked ADPR dissociates from the peptide, it appears to frequently take a portion of the Arg side chain with it. The result is a negative peptide remainder mass corresponding to the loss of the Arg reactive group that is both prevalent (66% of PSMs) and distinguishes ADPR on Arg from other residues. This is also reflected in the fragment remainder masses. The *b*- and *y*-ion series were found to consist of 40% and 26% ions shifted by -42, respectively (**Supplementary Data 5b**). Since only ions downstream of the modification site are expected to be shifted, we only expect to find half of all ions containing PTM-related mass shifts. The abundance of the Arg-specific fragment ions indicates that the modification itself should be easily localizable. We also found a noteworthy number of neutral loss-associated peptide remainder ions. When ADPR fragments after the primary ribose (**Fig 4b**, green), we would expect a peptide remainder mass of 114 Da if it were to remain intact. We do not find that mass, but instead find five sequential neutral losses of water from that mass. Equally of interest is that the neutral loss peaks—despite neutral losses not being unique to ADPRylated peptides—not found in unmodified spectra. Though counterintuitive, even common losses can produce PTM-specific peaks. By thinking of them as losses of almost the entire modification and a common neutral loss, it is easier to reconcile their uniqueness to specific modifications. In other words, a -17 peptide remainder mass (**Fig 4c**, red) will appear at the precursor *m/z* - 17 for unmodified peptides, but at precursor *m/z* – 558 for modified peptides.

### Use cases and applicability of diagnostic features

To investigate the extent to which co-fragmentation affects diagnostic feature characteristics, we leveraged our ability to identify large numbers of glycan diagnostic ions from the CPTAC IMAC-enriched dataset. This dataset represents 117 unique diagnostic ions, each found to be diagnostic for between 1 and 493 mass shifts, for a total of 13707 data points (**Supplementary Data 1**). Every diagnostic ion was evaluated individually for its ability to separate glyco and unmodified spectra based on its precision and AUC (**Supplementary Note 1**). This was repeated for the 64 unique peptide remainder masses observed between 1 and 344 times, totaling 2261 data points.

Here, precision can be interpreted as the probability ***y*** that a spectrum is a glyco spectrum given that the diagnostic ion is present in the spectrum at intensity ***x*** (**Fig 5a**). Diagnostic ion precision attenuates rapidly as the intensity increases, losing more than a third of its usefulness when it becomes the spectral base peak (average intensity 100.0). Because there is a detection limit for ions in mass spectrometers, less intense ions are also less likely to show up in spectra. As mentioned above, for co-fragmented spectra, the presence of the ions from the minor product is inversely proportional to the spectral purity and proportional to the ion’s intensity when its peptide is the major product. In other words, more intense diagnostic ions are more likely to appear in unmodified spectra because they can exceed the lower detection limit even for relatively pure unmodified PSMs. This is a trend can be reversed by taking intensity information into account rather than only checking for the presence or absence of the ion (**Fig 5b**). The AUC statistic here can be directly interpreted as the probability ***y*** that a diagnostic ion of intensity ***x*** drawn from a random modified PSM will be greater than the intensity of the same diagnostic ion drawn from a random unmodified PSM. After including intensity information, an ion’s ability to separate glyco and non-glyco spectra increases with intensity. Incorporating this feature into PTM-Shepherd allows us to detect diagnostic ions that are as ubiquitous as ADPR’s adenine ion, in 92.9% of off-target spectra in the HeLa dataset (**Supplementary Data 5a**). It also shows that researchers can effectively use intense diagnostic ions for scoring PTMs, but only if they empirically learn the distribution of intensities among unmodified PSMs beforehand.

**Figure 5:**
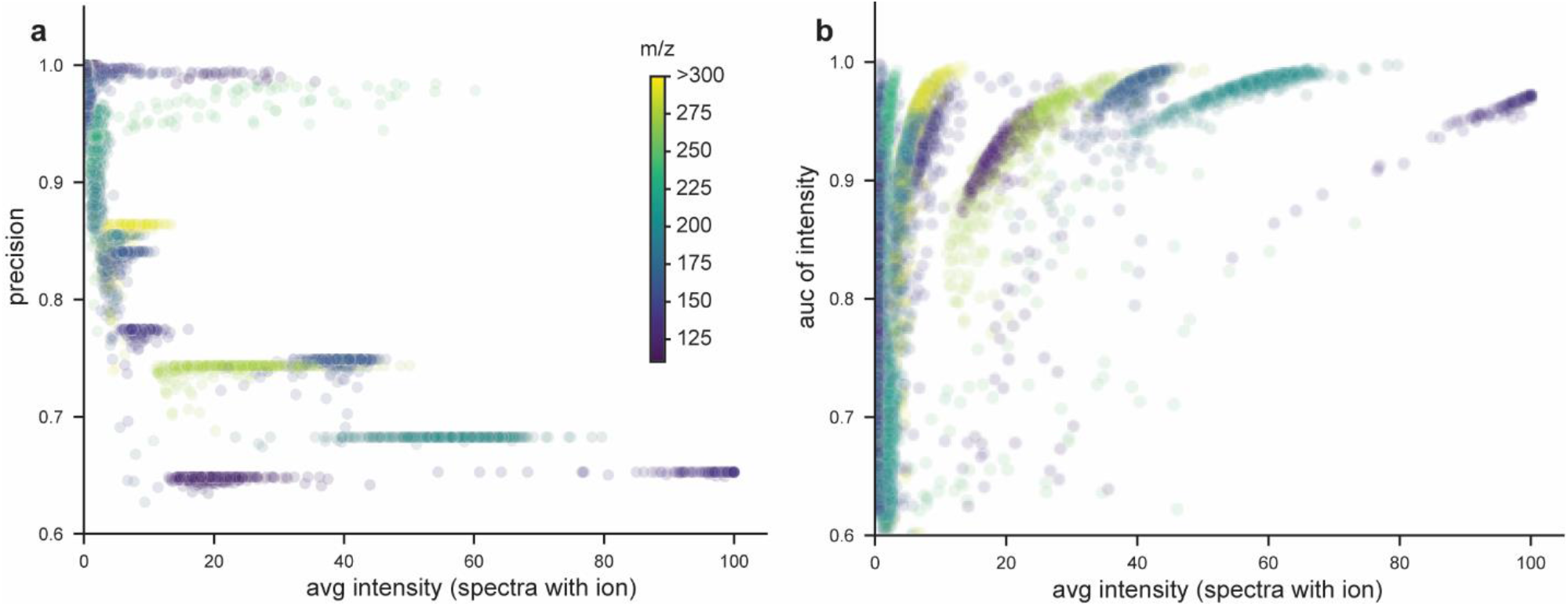
Trends in diagnostic remainder ions. (a) Relationship between diagnostic ions’ observed intensity and the precision of their presence. (b) Relationship between diagnostic ions’ observed intensity and the classification strength of their intensity as measured by AUC.

For peptide remainder masses, unlike diagnostic ions, precision does not attenuate with intensity (**Supplementary Fig 2**). As mentioned above, peptide remainder masses are mass-(although not sequence) specific. Co-fragmented peptides can only share peptide remainder masses if they share a mass that is indistinguishable at MS/MS mass accuracy, which is not guaranteed even for co-fragmented peptides with the same charge state. Excluding noise peaks that happen to fall within the tolerance of a theoretical peptide remainder ion, there should be few erroneously matched peptide remainder masses. The result is a very specific feature that does not attenuate as it gets more intense. Accordingly, peptide remainder ions discovered by PTM-Shepherd have many applications. Experiments performed with data-independent acquisition (DIA) have many co-fragmented peptides by design and present a prime opportunity for their use. Plus, with the advent of real-time searching, peptide remainder ions can also be used for instrumental enrichment^41^.

## Discussion

Our analyses show that PTM-Shepherd can be used to reliably identify diagnostic features for any modification of interest. In high-energy glycopeptide fragmentation, we showed that diagnostic ions for sialic acid could be identified without prior knowledge in a data-driven way, as well as finding two peptide remainder masses that had been described by experimentalists but neglected by cutting-edge glycopeptide search tools. In our discussion of a novel RNA-crosslinking workflow, we showed that we can easily automate experimental characterization in the FragPipe/PTM-Shepherd environment. Finally, our discussion of ADPR fragmentation demonstrated that fragmentation studies—traditionally done by hand with manual annotation of spectra or using custom tools^10^ and synthetic peptides^14^—could be automated and democratized to reach a broader audience and study PTMs without additional benchwork. We even found meaningful fragmentation patterns that would have been missed by annotation focused on modification structure alone. Although our analysis focused on demonstrating PTM-Shepherd’s capabilities, we also used our ability to generate diagnostic features in large numbers to better understand their nature. We showed that co-fragmentation of peptides presents a major issue for the precision of diagnostic ions in PTM analysis and explored ways to overcome it, as well as interrogating the utility of peptide and fragment remainder masses.

Automated diagnostic feature detection has wide-ranging applications across proteomics fields. Chemical probes can be characterized instantly, facilitating their development^19^. It could be advantageous to develop custom modification scores for localization-by-proxy strategies^42^ or as rescoring features in Percolator^43^. Furthermore, for enriched datasets or DIA-studies, the remainder masses identified by PTM-Shepherd might be the only reliable way to definitively identify labile modifications. There are myriad ways in which understanding modification behavior aids researchers, and thus we believe that the diagnostic feature detection enabled by PTM-Shepherd will be an invaluable tool in the analysis of proteomics data.

## Methods

### Spectral feature calculation

The first MS/MS spectral feature we analyze is raw spectral ions, such as immonium and oxonium ions, which we will refer to simply as diagnostic ions. All spectra from PSMs containing a given delta mass are stripped of matched *a-*ions, *b*-ions, and *y*-ions (by default). Spectra are also stripped of *a-, b-*, and *y-*ions that are found to be shifted by the PSM’s delta mass, preventing backbone fragments containing the modification from being counted as diagnostic ions. At this point, a spectrum can be thought of as a vector composed of ***m*** ions, where each ***ion***_***i***_ has a corresponding ***mz***_***i***_ and ***int***_***i***_ corresponding to the ion’s mass at charge state one and its intensity. All remaining ions are considered potential diagnostic ions and stored in a vector ***U*** of length ***m***. This can be represented as U = [(mz_1_, int_1_),⋯, (mz_m_, *int*_*m*_)].

The second MS/MS spectral feature we analyze in the MS/MS spectra is peptide remainder masses. All spectra from PSMs containing a given delta mass are stripped of shifted and unshifted *a-, b-*, and *y-*ions, as described above, before precursor remainder mass calculation. A theoretical peptide mass ***P*** of charge state one is calculated based on the peptide sequence and variable modifications identified for the PSM during spectral searching but excluding any MS1 mass shift. Then, the pairwise distance ***d*** between each remaining ion in the MS/MS spectrum and the theoretical peptide mass ***P*** is calculated and stored in a vector ***V*** of length ***m***, where ***m*** is the number of ions remaining in the spectrum after filtering. Each component ***V***_***i***_ contains the pairwise distance between ***P*** and ***mz***_***i***_ as well as the intensity ***int***_***i***_. This can be represented as V= [V_1_,⋯, V_*k*_] with each component V_*i*_ = (mz_*i*_ − P, int_*i*_). Intuitively, each component can be interpreted as what the precursor remainder mass and intensity would be if the ***i***th ion were a shifted precursor in the spectrum.

The third MS/MS spectral feature we analyze is fragment remainder masses. All spectra from PSMs containing a given delta mass are stripped of unshifted *a-, b-*, and *y-*ions only, allowing us to identify instances where the entire delta mass remains on the fragment ions. We reasoned that understanding how modifications affect individual ion series would provide insight into fragmentation patterns, so fragment remainder masses for *b-* and *y-*ions are calculated independently. For each fragment ion series, the peptide’s theoretical fragment ions of charge state one are calculated based on the peptide sequence and modifications identified for the PSM during spectral searching; the vector ***F*** holds each of ***n*** theoretical fragment ions, where ***n*** is the length of the peptide minus one and ***F***_***j***_ corresponds to the ***j***th fragment ion. Then, the pairwise distance between each remaining ion in the MS/MS spectrum and each theoretical fragment ion ***F***_***j***_ is calculated and stored in a matrix ***W*** of size ***m*** by ***n***, where ***m*** is the number of ions remaining in the spectrum. This can be represented as

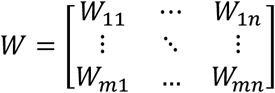

with each component W_*i*j_ = (*mz*_*i*_ − *F*_j_, *int*_*i*_). Intuitively, each matrix component can be interpreted as what the ***j***th fragment’s remainder mass and intensity would be if the ***i***th ion in the spectrum were the ***j***th theoretical fragment’s shifted counterpart.

### Identifying recurring features

We then determine which features represent the most intensity and are thus worthy of undergoing testing for enrichment. To do this, we place every value in a histogram with a bin width of 0.2 mDa spanning the range of possible features. For peptide and fragment remainder masses, the left tail of the histogram is truncated at -250 Da because values smaller than that would necessitate the losses of multiple residues. To account for uncertainty of ion position and smooth the histogram, the intensity of each ion is placed uniformly over an area equal to the MS/MS spectrum tolerance for the average ion in the histogram, i.e., the average inserted peptide or fragment mass. For diagnostic ions, a mass of 150 Da is used as the mass for smoothing. Furthermore, insertions into the histogram are normalized by the number of PSMs matching a particular peptide ion—that is, a grouping based on sequence, modification state, and precursor charge state—to prevent the inflation of features from abundant peptide ions.

Peaks in the histogram are defined by descending each side of a local maximum bin until a bin with zero intensity or a higher value is reached. Manual calibration found that a bin-to-bin tolerance of 1% was enough to prevent noisy bins from splitting peaks in two. Peaks are then integrated by summing histogram bins within the MS/MS tolerance without regard for adjacent peak boundaries. Any peak with an integrated area greater than 0.1% (by default), representing an average intensity greater than 0.1% of the base peak, is selected for downstream analysis. A final check is performed to remove redundant peaks where the least intense of any two histogram peaks that cannot be resolved under the provided MS/MS tolerance is removed.

### Identifying significant features

To find features specific to a particular mass shift, the full feature set—every major peak from the feature histograms above—needs to have features pruned from it that are not specific to the mass shift. We reasoned that peptides without mass shifts would be a good representative of a dataset’s noise, and as such testing whether features are more likely to appear among peptides with a particular mass shift than those without any mass shift would filter out non-modification-specific features.

Rather than using every PSM for what is inherently a noisy process, we select only those that are most likely to have the cleanest spectra. To do this, PSMs are first grouped by their peptide ion (sequence, modification state, and precursor charge state), then each group of PSMs has its lowest E-value representative selected for all downstream processing. The number of representative PSMs for each mass shifts is then capped at 1000 as we found that these gave reliable values across iterations. The 1000 representative PSMs are selected randomly, but internally a seed is provided for reproducibility.

Representative PSMs for every peptide ion with a particular mass shift and representative PSMs with no mass shift are first assembled, then every feature from the list of diagnostic ions, precursor remainder masses, and fragment remainder masses is quantified for each PSM in both lists. For spectra that do not have the diagnostic feature, the intensity is coded as a zero. Fragment remainder masses are likely to appear by chance solely based on the number of theoretical-to-experimental ion offsets calculated, so PSMs are considered to be missing a fragment remainder mass if there are fewer than two shifted ions of the ion type in the spectrum, i.e., fewer than two matching fragment remainder masses within feature matrix ***W*** for any ion type.

For every diagnostic feature tested, a series of metrics are produced for filtering noise peaks from real peaks. First, the lists of feature intensities from the unmodified and mass shifted PSMs are compared via a two-sided Mann-Whitney-U test with tie and continuity correction (adapted from the Hipparchus statistics library for Java, v1.8). E-values for each diagnostic feature are calculated by multiplying by the number of tests performed within the feature class for the current mass shift. By default, any feature with an E-value less than 0.05 is filtered out. A second metric to quantitatively assess the strength of the feature is included in PTM-Shepherd’s output: Area Under the Curve (AUC). This is commonly used as a measure of effect size for the Mann-Whitney U test and can be directly interpreted as the probability that a mass shifted PSM will have a higher intensity for this feature than an unmodified PSM. Second, we calculate a feature’s fold change of average intensity across all PSMs. Any features with fold change of less than 3.0 is filtered out by default. This metric primarily helps to identify diagnostic ions and non-specific but increased neutral losses for peptide and fragment remainders. Third, we filter out any features that are not sensitive for the modification, occurring in less than 25% of representative PSMs for diagnostic ions and peptide remainder masses. Owing to the multiple ion requirement for fragment remainder masses, this filter is reduced to 15% but is accompanied by an ion propensity filter required at least 12.5% of the identified ions within that series having the mass shift.

Fragment ions undergo an additional post-filtering processing step. Because a theoretical-experimental peak offset ***W***_***ij***_ is created for ***n*** theoretical ions in the theoretical ion series, a single peak in the experimental MS/MS spectrum produces a sequence specific pattern. For example, if the ***j***th residue produces an offset with fragment ***F***_***j***_ from the experimental ion ***i***, the same experimental ion responsible for that offset will also produce an “echo” offset from fragment ***F***_***j-1***_ equal to the original offset plus the mass of the residue at position ***j***. Similarly, it will produce an “echo” offset from fragment ***F***_***j+1***_ equal to the mass of residue ***j+1*** minus the original offset. Depending on the fragment ions containing the mass shift, some modifications can produce very weak signals for their primary mass shift but strong signals from shifted fragment ions upstream or downstream of the modification site. To correct for this, we check for residue enrichment both on and adjacent to the peptide site responsible for producing the mass shift. If any residue is found at position ***j+1*** for a modification more than 50% of the time, the fragment remainder mass is adjusted by subtracting that residue’s weight from the fragment remainder mass. If any residue is found at position ***j*** more than 50% of the time, the mass of residue ***j*** is added to the fragment remainder mass. With all fragments downstream of a peptide’s modification site carrying the mass shift, the residues responsible for these shifts should be roughly uniformly distributed across all 20 amino acids. Thus, any mass shift that is less prevalent than one of these adjusted offsets is unlikely to be a real peak, and reporting for fragment remainder masses is truncated after the first adjustment.

### Data processing

Four datasets were used throughout this manuscript. The first consists of a single Thermo Fisher Raw file “2021-2-23_EA_296_1A_Final.raw” from ProteomeXchange repository the PXD028853. This contains a cysteine chemoproteomic probe from Yan et al. (2022) that was used to demonstrate the algorithm. Data was collected on a Thermo Scientific Orbitrap Eclipse Tribrid in DDA mode and processed directly in FragPipe (v18.0) without conversion to mzML. Data was searched against the Uniprot reviewed protein sequences database retrieved on 13 June 2021 with decoys and common contaminants appended. An offset search was performed in MSFragger (v3.5)^5^ by loading the “Mass-Offset-Common-PTMs” workflow, replacing the offset list with 0 and 463.236554, and replacing the fixed cysteine carbamidomethylation with a variable one. “Write calibrated MGF”^24^ was turned on for the PTM-Shepherd^23^ diagnostic feature mining module, and “Diagnostic Feature Discovery” in PTM-Shepherd (v2.0.0) was enabled with default parameters. Filtering to 1% PSM, peptide, and protein levels was performed by Philosopher (v4.2.2)^47^.

The second dataset consists of the Clinical Proteomics Tumor Analysis Consortium (CPTAC) phosphorylation-enriched clear cell renal cell carcinoma (ccRCC) samples^28^ from the CPTAC data portal^44^. These 299 files represent TMT-labeled solid tumor or adjacent normal tissue from 110 human ccRCC patients. Samples were acquired on a Thermo Fisher Fusion Lumos in data-dependent acquisition (DDA) mode using high-collision dissociation (HCD). Thermo Fisher raw files were converted to mzML format using Proteowizard v3.0.11392^45^ with vendor peakpicking enabled. The 23 TMT-plexes were separated into separate experiment folders and processed using FragPipe v18.0. For the primary analysis, the default “glyco-N-TMT” workflow was used with minor changes to account for the goals of the analysis and experimental setup. Data was searched against the Uniprot reviewed protein sequences database retrieved on 13 June 2021 with decoys and common contaminants appended. During the MSFragger^5^ search, two variable phosphorylation modifications were allowed on the residues STY due to the expected enrichment of phosphorylated peptides and “Write calibrated MGF”^24^ was turned on for the PTM-Shepherd^23^ diagnostic feature mining module. In PTM-Shepherd, “Assign Glycans with FDR” was disabled, and “Diagnostic Feature Discovery” was enabled with default parameters. Finally, “Isobaric Labeling-Based Quantification” with TMT-Integrator was disabled. Filtering to 1% PSM, peptide, and protein levels was performed by Philosopher^47^. PTM-Shepherd was then run via command line to enable the reporting of isotopic peaks.

For the secondary analysis wherein known and discovered diagnostic ions were quantified, PTM-Shepherd’s “Diagnostic Feature Extraction” module was used with the ion list presented in Fig 2b. This was performed using the mzMLs rather than the deneutrallossed and deisotoped^24^ mgf files from MSFragger to prevent neutral losses that would be correlated under normal conditions from being anticorrelated in the analysis. For the tertiary analysis wherein the landscape of diagnostic features was explored, PTM-Shepherd was rerun, but with the filtering parameters for diagnostic ions and peptide ions set to 0 for “Min. % of spectra with ion” and 1 for “Min. intensity fold change.” PSM-level correlations for all PSMs with mass shifts >50 Da were computed for each diagnostic ion present in more than 50% of glycan mass shifts. Spectral ions are normalized to the base peak by default, creating nonlinear relationships between some ions and necessitating the use of Spearman’s rank correlation. Correlation was calculated using the Pandas package in Python.

The third dataset consists of a novel protocol for photoactivatable ribonucleoside-crosslinking from the ProteomeXchange repository PXD023401^25^. Only the two 4SU nucleotide-specific raw files from this repository were used. Samples were acquired on a Thermo Fisher Orbitrap Fusion Lumos using HCD fragmentation. Only the two 4SU-specific raw files from the repository were using in this analysis, and both samples were processing using FragPipe v18.0 directly without conversion to mzML. Samples were processed three times. The first, to find diagnostic features, was a standard open search using the FragPipe default “Open” workflow but with “Write calibrated MGF” and PTM-Shepherd’s “Diagnostic Feature Discovery” enabled with default settings. The second, to validate fragment remainder masses, was adapted from the default “Mass-Offset-CommonPTMs” workflow but with the mass offsets limited to 0, 226.0594, and 94.0168; “Labile modification search mode” enabled; “Y ion masses” and “Diagnostic fragment masses” removed; “Remainder masses” set to 94.0168 and 76.9903; “Write calibrated MGF” enabled; and PTM-Shepherd’s “Diagnostic Feature Discovery” enabled with default settings. The settings for the third analysis to validate an ammonium loss were identical to the second but without the 76.9903 fragment remainder mass. All analyses were run against the Uniprot database described above. Crystal-C^46^ was used to clean up open search results. Filtering to 1% PSM, peptide, and protein levels was performed by Philosopher^47^.

The fourth dataset consists of two samples from the ProteomeXchange repository PXD004245 corresponding to ADPR -enriched samples of mouse and HeLa origin^38^. The former is derived from mouse liver, processed in triplicate, and was acquired on a Thermo Fisher Orbitrap Q-Exactive Plus instrument in DDA mode using HCD. The latter was treated with H_2_O_2_ to induce oxidative stress, then collected in the same manner described above. Raw files were converted to mzML using Proteowizard v3.0.19296 with vendor peakpicking enabled. Both datasets were searched against their respective Uniprot reviewed sequence databased with decoys and common contaminants appended, with the mouse database retrieved on 27 September 2021 and the human database described above. Both datasets were searched separately in FragPipe v18.0 using the default “Labile_ADPR-ribosylation workflow with a few changes. During the MSFragger search, “Report mass shift as variable mod” was set to “No” so that PTM-Shepherd would register these ADPRs as mass shifts and “Write calibrated MGF” was enabled for the PTM-Shepherd diagnostic feature mining module. PeptideProphet^48^ and ProteinProphet defaults for “Offset search” were loaded, then PTM-Shepherd and its “Diagnostic Feature Discovery” Module were enabled.

## Supporting information

Supplementary Data Description

## Acknowledgements

This work was funded in part by National Institutes of Health grants (XXXAlexey). DJG was supported in part by the Proteogenomics of Cancer Training Program (5T32CA140044). We would like to thank Jong Woo Bae et al. for providing us early access to ProteomeXchange repository PXD023401. We would also like to thank our users for providing feedback on our tools.

## Contributions

DJG developed the algorithm and wrote the software, with DAP and FY assisting in the development of the algorithm and DAP assisting in the development of the software; DJG and AIN jointly analyzed the data and conceived the project while AIN supervised the project; DJG and DAP wrote the manuscript with input from all authors.

