## Supplementary Material for "Mining for ions: diagnostic feature detection in MS/MS spectra of post-translationally modified peptides"

Daniel J. Geiszler^1^, Daniel A. Polasky^2^, Fengchao Yu^2^, & Alexey I. Nesvizhskii^1,2,*^
^1^University of Michigan, Department of Computational Medicine and Bioinformatics

^2^University of Michigan, Department of Pathology

Supplementary Figures:

**Supplementary Figure 1:** Histograms of fragment remainders across mass shifts.

**Supplementary Figure 2:** Trends in peptide remainder ions.

Supplementary Tables:

**Supplementary Table 1:** Diagnostic features for a cysteine probe.

**Supplementary Table 2:** Supplementary Table 2: PTM-Shepherd diagnostic features for RNA Xlinking data from offset search.

Supplementary Notes:

**Supplementary Note 1:** Calculation of classification metrics from PTM-Shepherd

**Supplementary Note 2:** Analysis of ADP-Ribosylation ion specificity

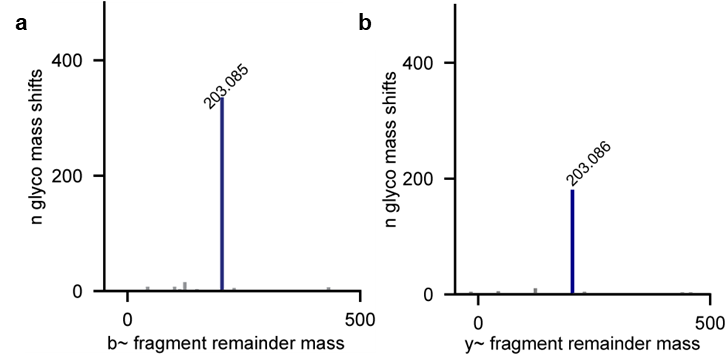

**Supplementary Figure 1: Histograms of fragment remainders across mass shifts.** (a) Fragments ions of *b-*type have a consistent and expected 203 Da remainder mass corresponding to a single HexNAc identified across many mass shifts. (b) Like *b-*ions, *y-*ions have a consistent and expected 203 Da remainder mass corresponding to a single HexNAc identified across many mass shifts.

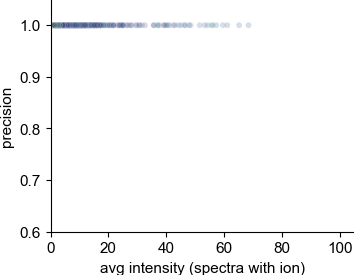

**Supplementary Figure 2:** **Trends in peptide remainder ions.** Unlike for diagnostic ions, peptide remainder ions retain their high precision due to being dependent on the mass of their host peptide and the lack of co-fragmentation interference.

| **peak apex** | | **mod**  **annotation** | **ion**  **type** | **mass** | **delta mod. mass** | **remainder propensity** | **percent PSMs**  **(mod)** | **percent PSMs**  **(unmod)** | **avg. intensity (mod)** | **avg. intensity (unmod)** | **intensity fold change** | **auc** |
| --- | --- | --- | --- | --- | --- | --- | --- | --- | --- | --- | --- | --- |
| 463.2364 | Unannotated mass-shift 463.2364 | | diagnostic | 301.1692 |  |  | 86.30 | 12.14 | 8.70 | 5.60 | 11.04 | 0.90 |
| 463.2364 | Unannotated mass-shift 463.2364 | | diagnostic | 327.185 |  |  | 88.80 | 20.32 | 8.83 | 5.76 | 6.70 | 0.89 |
| 463.2364 | Unannotated mass-shift 463.2364 | | diagnostic | 498.2314 |  |  | 98.80 | 51.98 | 31.51 | 14.87 | 4.03 | 0.89 |
| 463.2364 | Unannotated mass-shift 463.2364 | | diagnostic | 522.232 |  |  | 93.10 | 31.53 | 16.38 | 9.57 | 5.06 | 0.88 |
| 463.2364 | Unannotated mass-shift 463.2364 | | diagnostic | 424.2488 |  |  | 82.80 | 14.12 | 7.56 | 6.30 | 7.04 | 0.85 |
| 463.2364 | Unannotated mass-shift 463.2364 | | diagnostic | 227.084 |  |  | 85.40 | 22.3 | 5.81 | 4.34 | 5.12 | 0.85 |
| 463.2364 | Unannotated mass-shift 463.2364 | | diagnostic | 226.1012 |  |  | 45.70 | 2.51 | 2.34 | 1.97 | 21.60 | 0.72 |
| 463.2364 | Unannotated mass-shift 463.2364 | | diagnostic | 496.2164 |  |  | 46.10 | 7.39 | 3.60 | 6.65 | 3.38 | 0.69 |
| 463.2364 | Unannotated mass-shift 463.2364 | | diagnostic | 470.2254 |  |  | 40.40 | 3.17 | 2.64 | 6.70 | 5.03 | 0.68 |
| 463.2364 | Unannotated mass-shift 463.2364 | | peptide | 152.1038 | -311.133 |  | 49.70 | 0.00 | 12.98 | 0.00 | 100.00 | 0.75 |
| 463.2364 | Unannotated mass-shift 463.2364 | | peptide | 180.1032 | -283.133 |  | 47.50 | 0.00 | 13.94 | 0.00 | 100.00 | 0.74 |
| 463.2364 | Unannotated mass-shift 463.2364 | | b | 463.2364 | 0.00 | 44.11 | 48.50 | 0.53 | 33.32 | 28.37 | 100.00 | 0.74 |
| 463.2364 | Unannotated mass-shift 463.2364 | | b | 435.2420 | -27.9944 | 30.39 | 27.20 | 0.26 | 15.19 | 11.03 | 100.00 | 0.63 |
| 463.2364 | Unannotated mass-shift 463.2364 | | y | 463.2364 | 0.00 | 26.65 | 48.40 | 0.26 | 33.23 | 30.72 | 100.00 | 0.74 |

**Supplementary Table 1: Diagnostic features for a cysteine probe.** Remainder propensity scores are present only for *b-* and *y-* remainder masses. Features corresponding to the set of modified or unmodified PSMs used in comparisons are labeled as (mod) and (unmod), respectively. The difference between the MS1 mass shift and the observed fragment remainder masses (i.e., the lost mass) is enumerated in the “delta mod mass” column.

| **peak apex** | **mod annotation** | **ion** | **mass** | **delta mod. mass** | **remainder propensity** | **percent PSMs**  **(mod)** | **percent PSMs (unmod)** | **avg. intensity (mod)** | **avg. intensity (unmod)** | **intensity fold change** |
| --- | --- | --- | --- | --- | --- | --- | --- | --- | --- | --- |
| 226.0594 | Unannotated mass-shift 226.0594 | diagnostic | 133.0502 |  |  | 39 | 12.9 | 7.81 | 4.76 | 4.95 |
| 226.0594 | Unannotated mass-shift 226.0594 | diagnostic | 115.0396 |  |  | 31.1 | 10.4 | 6.04 | 3.84 | 4.70 |
| 226.0594 | Unannotated mass-shift 226.0594 | b | 94.0288 | 31.68 | -132.031 | 36.9 | 1 | 47.65 | 24.91 | 70.60 |
| 226.0594 | Unannotated mass-shift 226.0594 | b | 77.0052 | 18.47 | -149.054 | 17.9 | 1 | 29.43 | 13.13 | 40.13 |
| 226.0594 | Unannotated mass-shift 226.0594 | y | 94.0300 | 22.15 | -132.029 | 35.7 | 1 | 46.95 | 24.91 | 67.30 |
| 226.0594 | Unannotated mass-shift 226.0594 | y | 77.0068 | 15.11 | -149.053 | 17.4 | 0.9 | 28.81 | 13.05 | 42.68 |
| 94.0170 | Unannotated mass-shift 94.0170 | diagnostic | 215.0582 |  |  | 30.62 | 16.5 | 14.28 | 7.01 | 3.78 |
| 94.017 | Unannotated mass-shift 94.0170 | b | 94.0170 | 36.73 | 0 | 39.53 | 0.9 | 38.37 | 26.18 | 64.38 |
| 94.017 | Unannotated mass-shift 94.0170 | b | 77.0020 | 22.48 | -17.015 | 21.71 | 1 | 34.51 | 12.29 | 60.95 |
| 94.017 | Unannotated mass-shift 94.0170 | b | -19.0644 | 20.81 | -113.081 | 15.89 | 0.6 | 36.8 | 21.02 | 46.38 |
| 94.017 | Unannotated mass-shift 94.0170 | b | 66.0272 | 20.08 | -27.9898 | 15.89 | 1.3 | 19.82 | 23.58 | 10.28 |
| 94.017 | Unannotated mass-shift 94.0170 | y | 94.0170 | 23.75 | 0 | 39.53 | 0.9 | 38 | 26.18 | 63.76 |
| 94.017 | Unannotated mass-shift 94.0170 | y | 77.0064 | 14.41 | -17.0106 | 19.38 | 0.9 | 30.34 | 13.05 | 50.05 |

**Supplementary Table 2: PTM-Shepherd diagnostic features for RNA Xlinking data from offset search.** Remainder propensity scores are present only for *b-* and *y-* remainder masses. Features corresponding to the set of modified or unmodified PSMs used in comparisons are labeled as (mod) and (unmod), respectively. The difference between the MS1 mass shift and the observed fragment remainder masses (i.e., the lost mass) is enumerated in the “delta mod mass” column.

**Supplementary Note 1: Calculation of classification metrics from PTM-Shepherd.** PTM-Shepherd internally calculates a series of metrics to characterize diagnostic features. One of these is the Area Under the Curve of the Receiver Operating Characteristic Curve (AUC-ROC, or just AUC). This metric can be computed directly from the ***U***-statistic of the test group computed as part of the Mann-Whitney *U* test. The formula for this statistic is:

$${AUC}_{t}= \frac{U_{t}}{n_{t}n_{c}}$$

where ***t*** and ***c*** stand for test and control groups and ***n*** is the respective groups sample sizes. This metric has a useful interpretation as a rank probability statistic, i.e., it can be directly interpreted as the probability that a randomly chosen value from group ***t*** will be higher a randomly chosen value from group ***c***. Importantly, when calculated from the ***U***-statistic, AUC is not sensitive to class imbalances. This allows comparisons between diagnostic ions for mass shifts that have different numbers of PSMs, as in Fig 5a-c. The second metric calculated in the manuscript is precision, also known as positive predictive power. In classification problems, this metric is the inverse of FDR. Whereas FDR can be interpreted as the probability that a hit is a false positive given that it is positive, precision can be interpreted as the probability that a hit is a true positive given that it is positive. When using the presence of a particular feature to classify whether the spectrum contains the PTM of interest, this can be calculated easily from PTM-Shepherd output by the equation:

$$precision= \frac{{proportion}_{m}}{{proportion}_{u}+{proportion}_{m}}$$

where ***u*** and ***m*** correspond to unmodified and modified PSMs and ***proportion*** is the proportion of spectra containing the ion. Again, this metric is not sensitive to class imbalances when calculated this way, enabling direct comparisons between mass shifts with different numbers of PSMs.

**Supplementary Note 2: Analysis of ADP-Ribosylation ion specificity.** We attempted to do searches for ADP-Ribosylated (ADPR) peptides using parameters as close to FragPipe defaults as possible to make them reproducible for other users. The default ADPR workflow includes both a labile and variable modification search for ADPR, allowing competition between fragmented and intact forms of the modification. Because the zero bin is defined as peptides with no mass shift and PTMs identified as variable mods have no mass shift, some ADPR-containing peptides are also present in the unmodified peptide bin. However, these only account for roughly 1 in 7 PSMs in the unmodified bin, and as such cannot be driving the trends discussed here.
