## Supplementary Data Description for "Mining for ions: diagnostic feature detection in MS/MS spectra of post-translationally modified peptides"

**Title:** Supplementary Data 1

**Description:** Three tables from PTM-Shepherd analysis of RNA crosslinking data. a) PTM-Shepherd mass shift profile of RNA Xlinking data. The PTM-Shepherd global.profile.tsv table for this analysis was created using the default diagnostic feature mining workflow. Output was used to determine which mass shifts correspond to the intact and fragmented RNA moiety. b) PTM-Shepherd diagnostic features for RNA Xlinking data from open search. The PTM-Shepherd global.diagmine.tsv table for this analysis was created using the default diagnostic feature mining workflow. Output was used to determine which features would be included in a second pass offset search. Used to guide analyses for Figure 2.

**Title:** Supplementary Data 2

**Description:** PTM-Shepherd diagnostic features (global.diagmine.tsv) for analysis of IMAC-enriched CCRCC data. Used in the creation of Figures 3a, c, Supplementary Figures 1 and 2, and Figure 5.

**Title:** Supplementary Data 3

**Description:** Spearman correlation matrix between ion intensities across spectra for the analysis of IMAC-enriched CCRCC data. Used in the creation of Figure 2b.

**Title**: Supplementary Data 4

**Description:** Two tables (global.diagmine.tsv) from PTM-Shepherd analysis of ADP-ribosylation. a) PTM-Shepherd diagnostic features for ADPR from mouse. Used in the creation of Figures 4a, c, d. b) PTM-Shepherd diagnostic features for ADPR from HeLa. Used in the creation of Figures 4a, c, d.

**Table Column Guide**

**global.profile.tsv**

global.profile.tsv reports the most prominent features from PTM-Shepherd analysis of mass shifts observed from FDR-filtered open search results. Each row corresponds to a different detected mass shift, thus not all PSMs will be represented in this table. Please note that mass shifts are annotated based on UniMod mapping, thus they are not definitive chemical identities and should be used as a starting point along with localization and amino acid enrichment information. Unless otherwise indicated, values are summed from all datasets in the analysis. Column contents are listed below.

- **peak_apex** apex of the detected delta mass peak (in Da)
- **peak_lower** lower bound of the detected peak (Da), determined by precursor tolerance or the detection of an adjacent peak
- **peak_upper** upper bound of the detected peak (Da), determined by precursor tolerance or the detection of an adjacent peak
- **PSMs** the number of PSMs contained within the peak boundary (bin), reported for each dataset if multiple datasets are used as input
- **peak_signal** relative measure of peak prominence/quality. In noisy regions of the delta mass histogram, values are penalized
- **percent_also_in_unmodified** the percentage of PSMs in this mass bin with a corresponding PSM in the unmodified bin
- **mapped_mass_1** primary modification annotation derived from Unimod, all isobaric modifications listed and separated by “/”
- **mapped_mass_2** if the delta mass peak is a combination of two masses, a second modification annotation is listed here. As with mapped_mass_1, all isobaric modifications are listed and separated by “/”
- **similarity** MS/MS spectral similarity of modified peptides compared to their unmodified counterparts. When multiple modified-unmodified comparisons are done for a single peptide, these cosine similarity scores are averaged for the peptide. The peptide scores are then averaged across all peptides in the mass shift bin. These comparisons are only done for peptides of the same charge state.
- **rt_shift** retention time shift comparing modified peptides to their unmodified counterparts. When multiple modified-unmodified comparisons are done for a single peptide, the retention time shifts are averaged for the peptide. The peptide shifts are then averaged across all peptides in the mass shift bin. Individual comparisons are only done for peptides in the same LC-MS run. Units are usually seconds but can vary by instrument type
- **int_log2fc** log2 fold-change of average intensity for matched shifted/unshifted peptides, computed as described above. Peptides affect by sample preparation artifacts tend to be lower abundance than their unshifted counterparts, thus this value will be low in these cases
- **localized_PSMs** number of PSMs for this delta mass that showed at least one additional matched ion when the mass shift is placed on a residue
- **n-term_localization_rate** percentage of PSMs with an uninterrupted string of localized residues from the N-terminus. This is calculated differently from other enrichment scores due to the difference in assumptions underlying N-terminal and residue-specific localization, so these values cannot be directly compared to the amino acid enrichment scores.
- **AA1** amino acid/residue most enriched (most likely to harbor the mass shift) compared to other residues
- **AA1_enrichment_score** equivalent to the odds the delta mass is localized to AA1 compared to other residues
- **AA1_psm_count** weighted number of PSMs where the mass shift localized to AA1. Shifts localizing to multiple residues are divided by the number of localized residues in the spectra, so this is an estimated number of PSMs localized to a particular residue
- (same enrichment_score and psm_count columns for AA2 and AA3 if multiple amino acids are likely to harbor the mass shift)
- **[experiment]_PSMs** number of PSMs with a mass shift in this bin
- **[experiment]_percent_PSMs** number of PSMs from the previous column as a percentage of total PSMs
- **[experiment]_peptides** number of unique peptide sequences with a mass shift in this bin
- **[experiment]_percent_also_in_unmodified** percentage of peptide sequences with a mass shift in this bin that are also found in the zero mass shift bin

**global.diagmine.tsv**

global.diagmine.tsv is a mass shift-centric table that contains the diagnostic features identified for every mass shift. Please note that only mass shifts with diagnostic features detected are reported in the table. Contents of each column are listed below.

- **peak_apex** This field contains the apex of the detected MS1 peak (Da) present in the global.profile.tsv file from PTM-Shepherd.
- **mod_annotation** This field contains the mass shift annotations present in the global.profile.tsv file from PTM-Shepherd. When a mass shift is found to be the combination of two mass shifts, the “Potential Modification 1” and “Potential Modification 2” columns are merged with a semicolon.
- **type** This field can take one of several values. “diagnostic” refers to diagnostic ions, the ions that can be located directly in the spectrum. “peptide” refers to peptide remainder masses, mass shifts that indicate an ion’s presence at a particular distance from an unshifted peptide. Six other values are possible based on parameter setting, each corresponding to one of the major ion series.
- **mass** This field contains the mass of the diagnostic feature. Peptide and fragment remainder masses will have the mass shift away from the theoretical ion. Diagnostic ions will have the m/z of the observed ion, so a non-neutral mass.
- **delta_mod_mass** This field contains the mass that was lost from the original mass shift to arrive at the remainder mass. (Note: only present for peptide and fragment remainder masses.)
- **remainder_propensity** This field contains the average percentage of ions from a particular series that are shifted. For example, a peptide capable of producing 10 *b*-ions with 2 ions identified ions shifted by the remainder mass and 2 identified ions unshifted would have a propensity of 50%. The propensity score for every representative PSM within a mass shift bin is averaged. (Note: only present for fragment remainder masses.)
- **percent_mod** This field contains the percentage of representative mass shifted PSMs that contain the ion at any intensity.
- **percent_unmod** This field contains the percentage of representative unshifted PSMs that contain the ion at any intensity.
- **avg_intensity_mod** This field contains the average intensity of the ion among representative mass shifted PSMs where the ion is present. To calculate the average across all representative mass shifted spectra, calculate (avg_intensity_mod * percent_mod / 100). Because multiple ions can be matched for fragment remainder ions, this contains the average of the summed intensity of matched ions for each representative PSM.
- **avg_intensity_unmod** This field contains the average intensity of the ions among representative unshifted PSMs where the ion is present. To calculate the average across all representative mass shifted spectra, calculate (avg_intensity_mod * percent_mod / 100). Because multiple ions can be matched for fragment remainder ions, this contains the average of the summed intensity of matched ions for each representative PSM.
- **intensity_fold_change** This field contains the fold change in intensity when comparing the modified to unmodified peptides. This uses intensity across all spectra and can be calculated via (avg_intensity_mod * percent_mod) / (avg_intensity_unmod * percent_unmod).
- **auc** This column contains the AUC-ROC statistic for the intensity-based classification of this ion. It is calculated from the U statistic from the Mann-Whitney U Test. This statistic adjusts the two groups such that they are assumed to be of equal size.
